# Chromatin interaction data visualization in the WashU Epigenome Browser

**DOI:** 10.1101/239368

**Authors:** Daofeng Li, Silas Hsu, Deepak Purushotham, Ting Wang

**Affiliations:** Department of Genetics, Edison Family Center for Genome Sciences and Systems Biology, Washington University School of Medicine, St. Louis, MO, USA

## Abstract

**Motivation:** Long-range chromatin interactions are critical for gene regulations and genome maintenance. HiC and Cool are the two most common data formats used by the community, including the 4D Nucleome Consortium (4DN), to represent chromatin interaction data from a variety of chromatin conformation capture experiments, and specialized tools were developed for their analysis, visualization, and conversion. However, there does not exist a tool that can support visualization of both data formats simultaneously.

**Results:** The WashU Epigenome Browser has integrated both HiC and Cool data formats into its visualization platform. Investigators can seamlessly explore chromatin interaction data regardless of their underlying data format. For developers it is straightforward to benchmark the differences in rendering speed and computational resource usage between the two data formats.

**Availability:** http://epigenomegateway.wustl.edu/browser/.

## 1 Introduction

The WashU Epigenome Browser (Zhou, et al., 2011) has been serving as an integrated data visualization and analysis platform for the epigenomics research community worldwide since 2011. Since its inception, the Browser has undergone continuous development, adding new visualizations, data sets, functions, and new features. The Browser currently hosts over 50,000 datasets from international consortiums including Roadmap Epigenomics (Roadmap Epigenomics, et al., 2015), ENCODE (Encode Project, 2012) and IHEC (Stunnenberg, et al., 2016) projects, and continues to incorporate 4DN data as the consortium scales up its data production (Dekker, et al., 2017).

One of the primary goals of the WashU 4DN Data Center is to develop solutions that insightfully display 4DN genomics data, which examine the highly organized and three-dimensional structure of eukaryotic chromosomes. The WashU Epigenome Browser has been an important tool for exploring how eukaryotic genomes function as nonlinear systems since the 2013 invention of new long-range interaction tracks for Hi-C and ChIA-PET data (Zhou, et al., 2013). The rapid development of genomic technologies and bioinformatics tools for interrogating chromatin interaction data has propelled two data formats to become the top choice for representing Hi-C data, i.e., .hic format and .cool format, .hic was invented in 2016 (Durand, et al., 2016). Most Hi-C data in the public domain has been processed into .hic format by juicer (Durand, et al., 2016). Correspondingly, Juicebox is perhaps the most popular tool for .hic data visualization. In contrast, .cool was invented in 2016 [https://github.com/mimylab/cooler], and HiGlass (Kerpedjiev, et al., 2017) is a newly developed visualization platform for .cool datasets. While both .hic and .cool formats store genomic contact matrices as highly compressed binary files that allow fast random access, they differ in several important ways. For example, .cool format is based on the standardized HDF5 format; hic is also a compressed binary format supporting random data access, but it is not based on an existing standard and tends to have a larger footprint than .cool (Figure 1A). The 4DN consortium uses both .hic and .cool file formats to represent high-throughput chromatin interaction by Hi-C. Although tools for converting between the two data formats exist [https://github.com/4dn-dcic/hic2cool], there isn’t a system in which both data formats co-exist, especially when it comes to their visualization which is where investigators begin to engage with these type of data.

**Figure 1.**
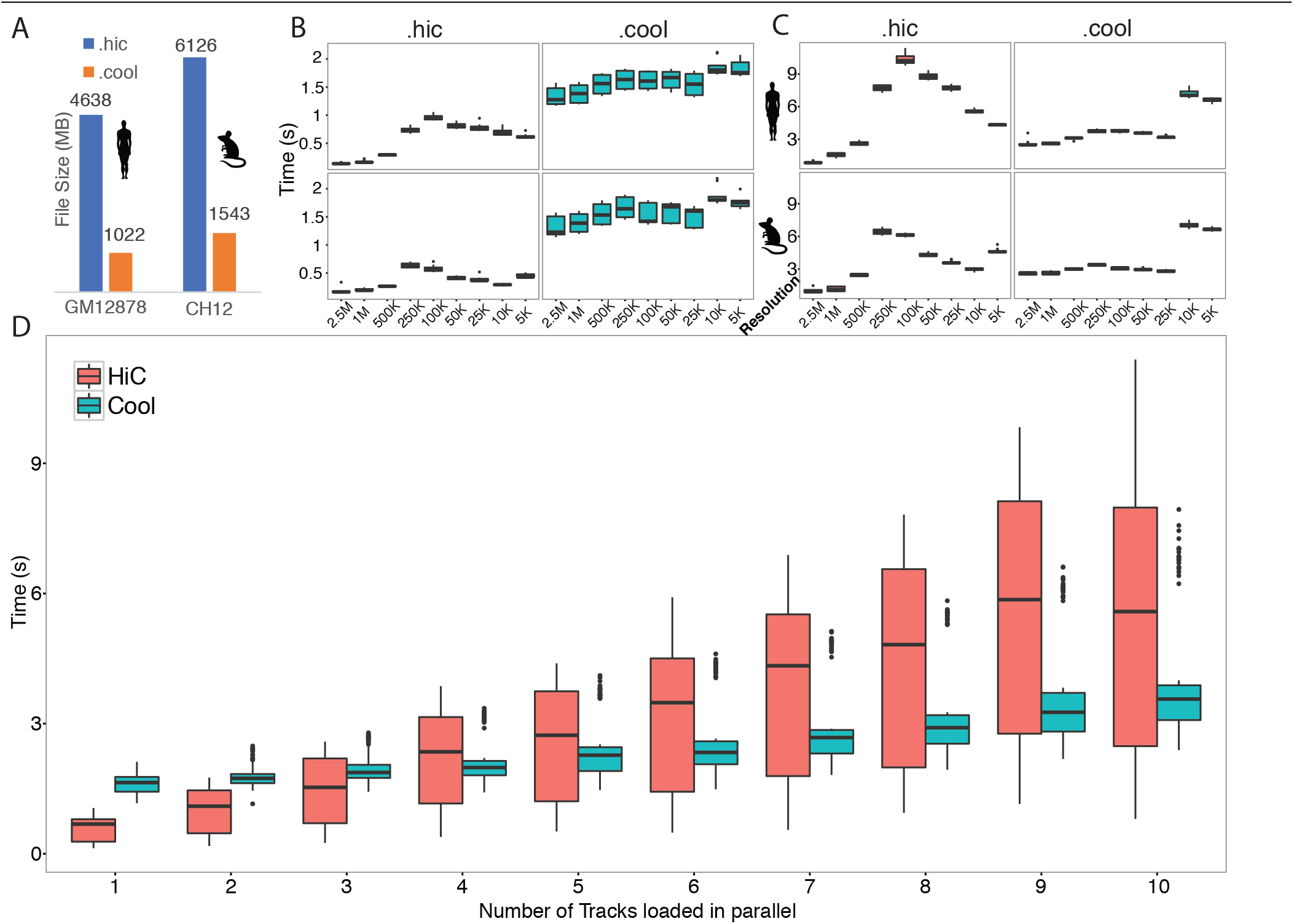
benchmark of hie and cool format. A. Comparison between file sizes of this 2 file formats. B. Data fetching speed for single track of hic and cool format in human and mouse data. C. Data fetching speed for 10 tracks in parallel. D. Relationship between data fetching speed versus number of tracks loaded in parallel.

Recently, the WashU Epigenome Brower has significantly expanded its capacity in visualizing long range chromatin interaction data. In addition to displaying tab-delimited text files representing genome interactions, the Browser now supports simultaneous visualization of .hic data and .cool data, liberating investigators from having to pick and choose between tools and data formats, especially when an investigator wishes to compare and explore their own Hi-C data and publicly available chromatin interaction data. For users who are interested in comparing performance between these two advanced data formats, the comparison is straightforward since they co-exist on the same platform. In this report, we present our Browser’s ability to visualize data in both .hic and .cool formats, as well as exemplar benchmarking that compares these two formats.

## 2 Methods

We implemented both .hic and .cool visualization tracks at WashU Epigenome Browser.

We timed and compared the tracks’ rendering speed at various resolutions for two datasets:

- Human GM12878, B-cell lymphoma (https://www.ncbi.nlm.nih.gov/geo/query/acc.cgi?acc=GSM1_551551)
- Mouse CH12, B-cell lymphoma, analog to human GM12878: (https://www.ncbi.nlm.nih.gov/geo/query/acc.cgi?acc=GSE6_3525)

The original data was available in .hic format. We used a utility called hic2cool (https://github.com/4dn-dcic/hic2cool) to convert the .hic files to the .cool format.

For each dataset, we fetched data from 1 Mb region. We also tested up to 10 parallel queries.

The test was done in Google Chrome (Version 60.0.3112.101 (Official Build) (64-bit)), on a 2015 iMac running MacOS Sierra version 10.12.5 with 3.3 GHz Intel Core i5 processor.

## 3 Results

The Browser can now display multiple formats of the same long-range interaction data side-by-side, quickly, and accurately (Supplemental Figure 1). The patterns displayed by the WashU Browser faithfully recapitulates published images [e.g., Figure 2A from (Sanborn, et al., 2015)], as well as images displayed by Juicebox and HiGlass (Supplemental Figure 1).

Our benchmark results suggest that rendering .hic data is faster than .cool data for a single track, but .cool is faster than .hic when displaying more than 3 tracks. The most plausible explanation is that .hic files are being read entirely and sequentially by client-side JavaScript code, which runs on a single thread. In contrast, Cool files are processed on server-side, which leverages the multicore capabilities of the server. This comparison immediately reveals future directions for performance improvement.

### 3.1 Implementations

We took advantage of existing tools that handle .hic and .cool files. To display the .hic format, we adopted the juicebox.js library which provides easy reading of .hic data via the Internet, and modified the API code to suit the needs of the Browser. The current architecture of HDF5 format does not allow remote access; thus, we processed the data using Cooler and hosted the data server-side, and implemented a REST API from which clients can request data.

### 3.2 Benchmarking

For the same data, .cool files took roughly 25% of the space that .hic files use (Figure 1A). On the client side, data fetching speed of .hic file was faster than cool file when querying a single track (Figure 1B), while .hic was slower than .cool when 10 tracks were queried in parallel (Figure 1C). Data fetching time of .hic data increased more drastically than .cool data as the number of tracks increased, as illustrated in Figure 1D. We think the reason is that .hic track visualization depends on juicebox.js, a client side JavaScript library; thus, the client incurs a high computational burden when a large number of tracks are loaded. This is confirmed by our CPU and RAM usage comparison (Supplemental Table 1). In contrast, for the .cool format, the client side only downloads data while the server handles the main computational burden for data access; thus, the time spent on the client side only increases slightly when many tracks are loaded in parallel.

Note that since we disabled the web browser cache during benchmarking, the benchmarks best simulate the performance of adding new tracks and the initial render. In the real world, data fetch and render will be faster after the initial load, especially when repeatedly requesting the same general region.

### 3.3 Data hub

The Browser has already had public .hic data hubs available for some time, which consist of tracks from Juicebox (206 tracks for human and 74 for mouse genome). For better serving the research community, we have organized .cool data hubs for the human and mouse genomes, and have populated the hubs with tracks converted from our .hic hubs. As of this writing, our public hubs contain 185 .cool tracks for the human genome, and 79 for the mouse genome. Supplemental tables 2 and 3 contain additional details, including the cell lines and publications from which the tracks originate. As the research community generates more data, we will be able to expand the public hubs even more.

## Acknowledgements

We thank Jim Robinson for developing juiceboxjs; we thank the Mirny lab for developing the Cooler package.

